# HuluFISH non-denaturing in situ detection of genomic DNA opened by CRISPR-Cas9 Nickase and Exonuclease

**DOI:** 10.1101/2021.12.23.473974

**Authors:** Kang Han, Sheng Liu, Yongsheng Cheng

## Abstract

DNA fluorescence in situ hybridization (FISH) has been widely used in diagnosis and genetic research. Traditional Bacterial artificial chromosome (BAC) or oligonucleotide based probe to detect DNA in situ is only effective when the target is relatively large, usually over 150Kb DNA fragments. And it involves heat denaturation steps to open the DNA for in situ hybridization. The heat process can affect the fine structure of nuclei. Here we reported a novel method based on Cas9 nickase and exonuclease digestion of double strand DNA and permanently mark the DNA in single strand state for FISH. With this novel design, we detected non-repetitive genomic loci as small as 2Kb.

## Introduction

DNA FISH is a powerful technique in detecting chromosomal structures and has been widely used in study chromatic structure and clinical diagnosis of chromosomal abnormalities. However, the DNA FISH needs the genomic DNA to be opened by heat denaturation and relatively long target region to generate sufficient hybridization signal for detection. The usual BAC (Bacterial Artificial Chromosome) based DNA FISH probe spans dozens of kilobases (kb) or even hundreds of kb in genomic DNA. For fine chromatic structure study, the heat denaturation and long target region sometimes are prohibitive, especially when the target region is less than 10 kb like an exogenous gene introduced into the genome in cell and gene therapy.

The CRISPR-Cas9 (Streptococcus pyogenes) system has been extensively explored in studying genome structure both in vivo and in vitro (Chen et al., 2013; Deng et al., 2015; Ishii et al., 2019; Knight et al., 2015; Wang et al., 2021). The CRISPR-Cas9-sgRNA ribonucleoprotein complex can target a specific loci on genomic and label or open genome for hybridization.

Here we are presenting a novel method (CRISPR-ExoFISH) based on the CRISPR-Cas9 system to open the genome loci and digest the specific loci to be single strand DNA (ssDNA) for permanent accessibility for DNA FISH probes (Figure 1). Once the DNA is opened by CRISPR-Cas9 (H840A mutant) on the target strand (TS), a T7 exonuclease starts to digest the non-targeting strand (NTS) of the CRISPR-Cas9 system and leave the targeting strand intact for hybridization. For the following FISH probe hybridization, heat denaturation is not needed and a shorter region below 5kb is sufficient for the DNA FISH probe to bind and generate specific signals.

**Figure 1.**
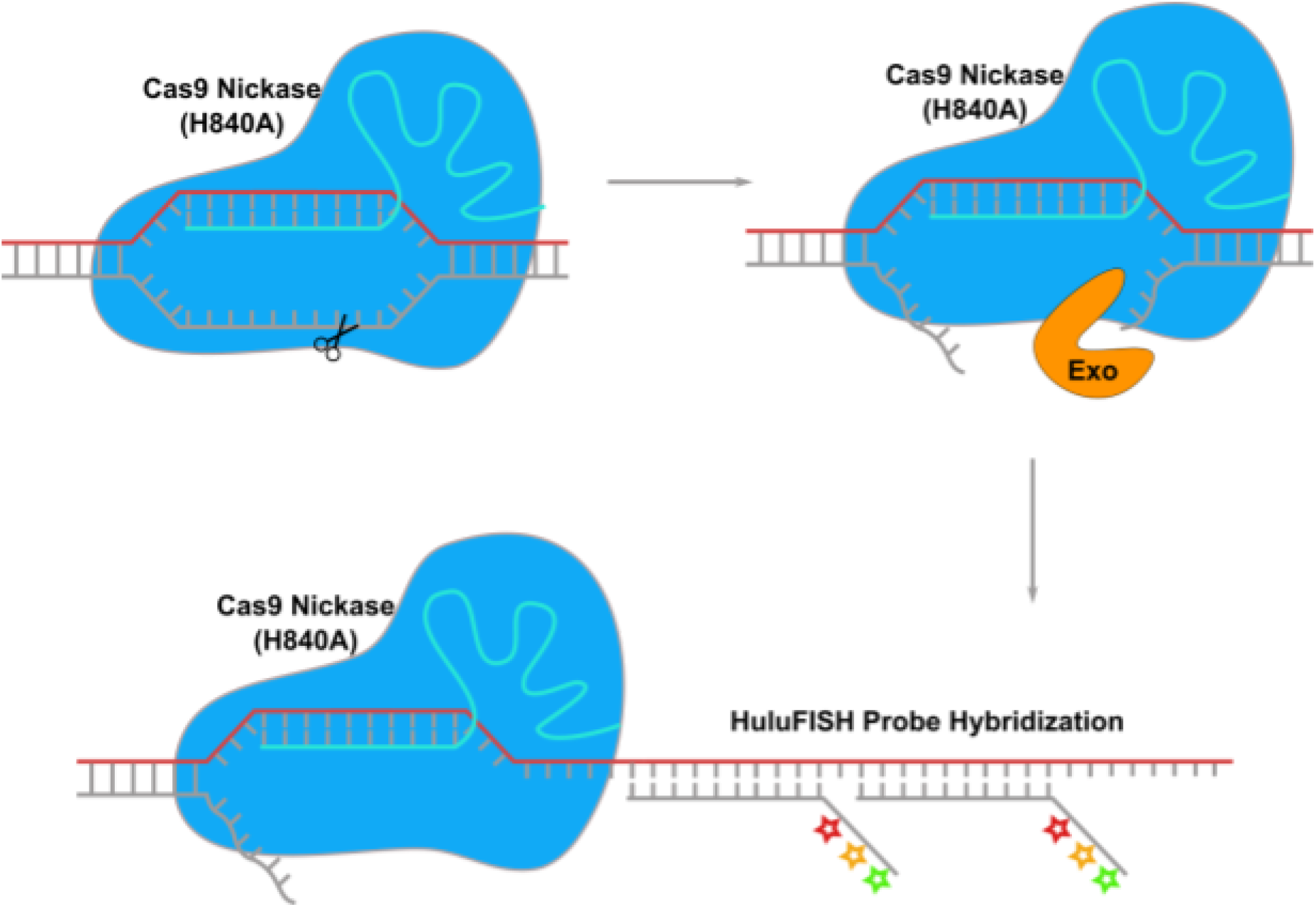
CRlSPR-ExoFISH workflow. Genomic DNA loci is binded by Cas9 nickase (H840A)-sgRNA (cyan) ribonucleoprotein complex and cut at NTS strand (grey). Exonuclease find the flapped DNA on NTS strand and digest the NTS stand. Then HuluFlSH probe is hybridizing to the TS stand (red).

## Method and Materials

### sgRNA synthesis by in vitro transcription (IVT)

sgRNA was produced with the ENGen SgRNA synthesis kit (NEB, E3322). sgRNA guide sequences were designed by customized script at PixelBiotech GmbH for Human Telomere, alpha-satellite DNA (ASAT), and the non-repetitive region of MUC4 gene (Intron 1, 2kb long target). Target-specific DNA oligo templates were designed according to the protocol for ENGen SgRNA synthesis kit. The sgRNA IVT was following the protocol from the kit. For multiple sgRNA synthesis in one reaction, the total concentration of oligo DNA template was adjusted to 250nM final concentration. The sgRNA was purified with the Monarch RNA Cleanup kit (NEB, T2040).

### CRISPR-ExoFISH

Hela cells were fixed by cold methanol:acetic (3:1, stored at −20°C) for 10 min and stored at −20C or immediately used. RNase A (NEB, T3018) treatment at 0.1mg/ml in the 1xPBS for 30 min at 37°C is optional to remove background staining from cellular RNA. After RNase A treatment, the cells were washed with 1xPBS at 65C for 10 min each, 3 times. Cas9-sgRNA ribonucleoprotein complex was reconstituted in binding buffer (20mM HEPES, pH 7.5, 100mM KCl, 5mM MgCl2, 5% (v/v) Glycerol, 0.1% (v/v) Tween-20, 1% (w/v) Bovine serum albumin (BSA), 1mM Dithiothreitol (DTT), 0.4U/ul RNase inhibitor Murine (NEB, M0314)) at the concentration of 200nM each (1:1 ratio for Cas9:sgRNA concentration) for 10 min at room temperature. Fixed cells were washed with the binding buffer for 10 min at room temperature. Then the cells were incubated with 200nM Cas9-sgRNA ribonucleoprotein complex in the binding buffer with 1U/ul T7 exonuclease (NEB, M0263) for 1 hour at 37°C. Then the cells were washed with 1xPBS for 5 min. After washing with 1xPBS for 3 times in 5 min, cells were hybridized with a 320nM HuluFISH probe in HuluHyb (PixelBiotech GmbH, PB102) for 1 hours at 37°C. After hybridization, the cells were washed with 2xSSC, 20% formamide at 37°C for 10 min for two times, and then washed with 1xPBS for 5 min before mounting in the VectorShield HardSet (Vector Laboratories, H-1400). Stained cells were imaged under Leica TCS SP8 confocal microscope.

## Results

We tested CRISPR-ExoFISH on 3 targets in human cells. In Figure 2A-B, telomere sgRNA were designed against the repeat TAAGGG in Human telomeres (sequence details in table 1). Without sgRNA and Cas9, HuluFISH probes for telomeres could not bind it all onto the genome without heat denaturation. In the facilitation of Cas9-sgRNA complex, HuluFISH probes efficiently hybridize to the telomere region. This indicates the ssDNA nature of telomeres after Cas9-sgRNA binding and T7 exonuclease digestion. Another repeat region in the genome, alpha satellite DNA (ASAT) was chosen for further validation of this novel method (Figure 2C-D). All repeated genomic regions show specific staining compared to the negative control without Cas9-sgRNA opening and digesting the genomic DNA.

**Figure 2.**
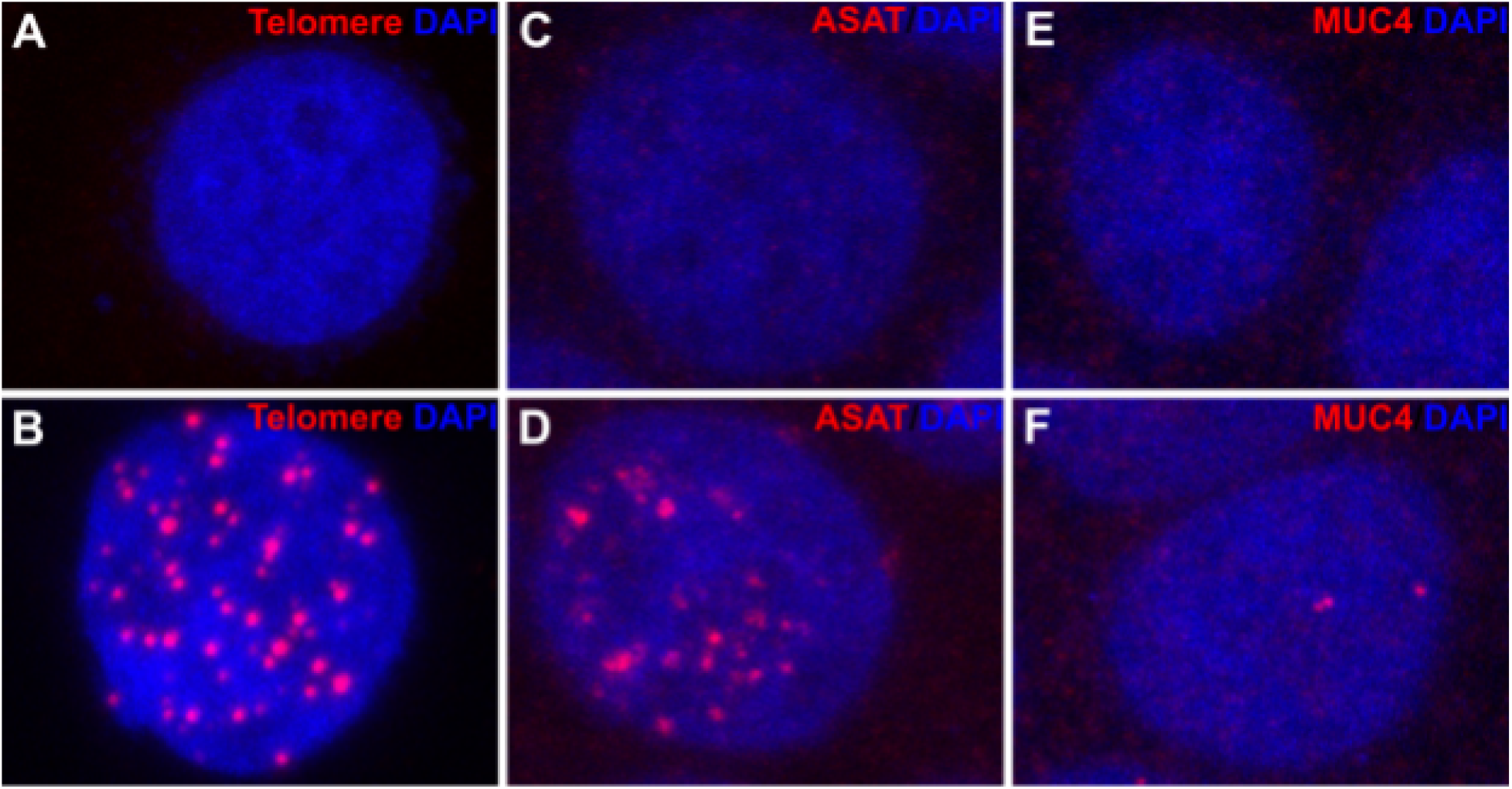
CRlSPR-ExoFISH of Human genomic targets. (A, C, E) Hela cell stained with HuluFISH probe for telomere, ASAT, and MUC4 respectively, (B. D, F) Hela cell were fixed and stained with HuluFISH probe against Human telomere, ASAT and MUC4 respectively, after genomic DNA of telomere, ASAT and MUC4 is opened and digested as ssDNA.

Despite the repeat region, CRISPR-ExoFISH was also employed to detect non-repetitive genomic loci. MUC4 intron 1 was chosen for the validation (Figure 2E-F). A 2kb region was selected for sgRNA design and 9 sgRNAs were selected for in vitro transcription and assembly with Cas9. Additionally 63 HuluFISH probes were designed for the 2kb region in MUC4. The results showed specific staining of the MUC4 gene intron 1.

## Appendix 1

**Table 1.**
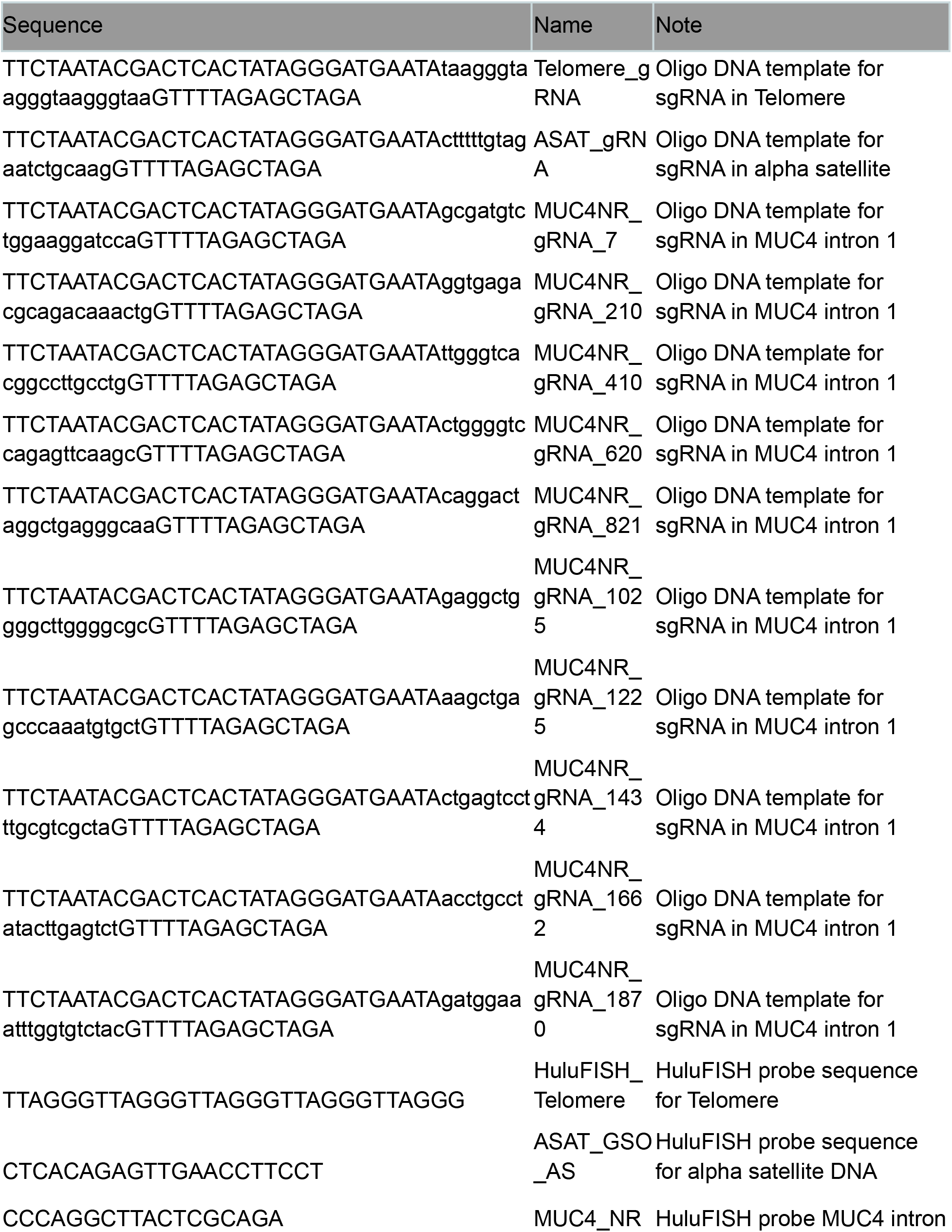
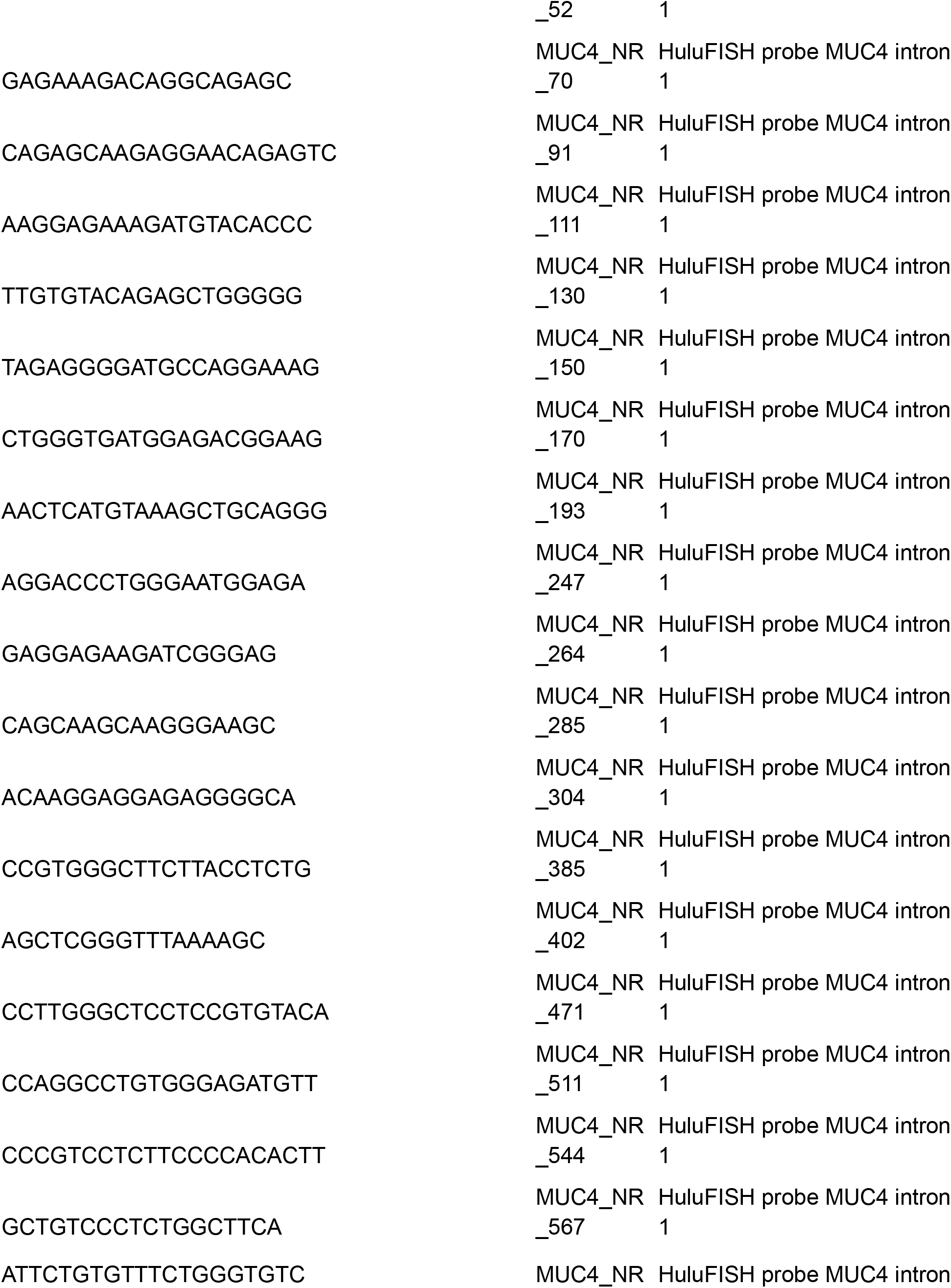
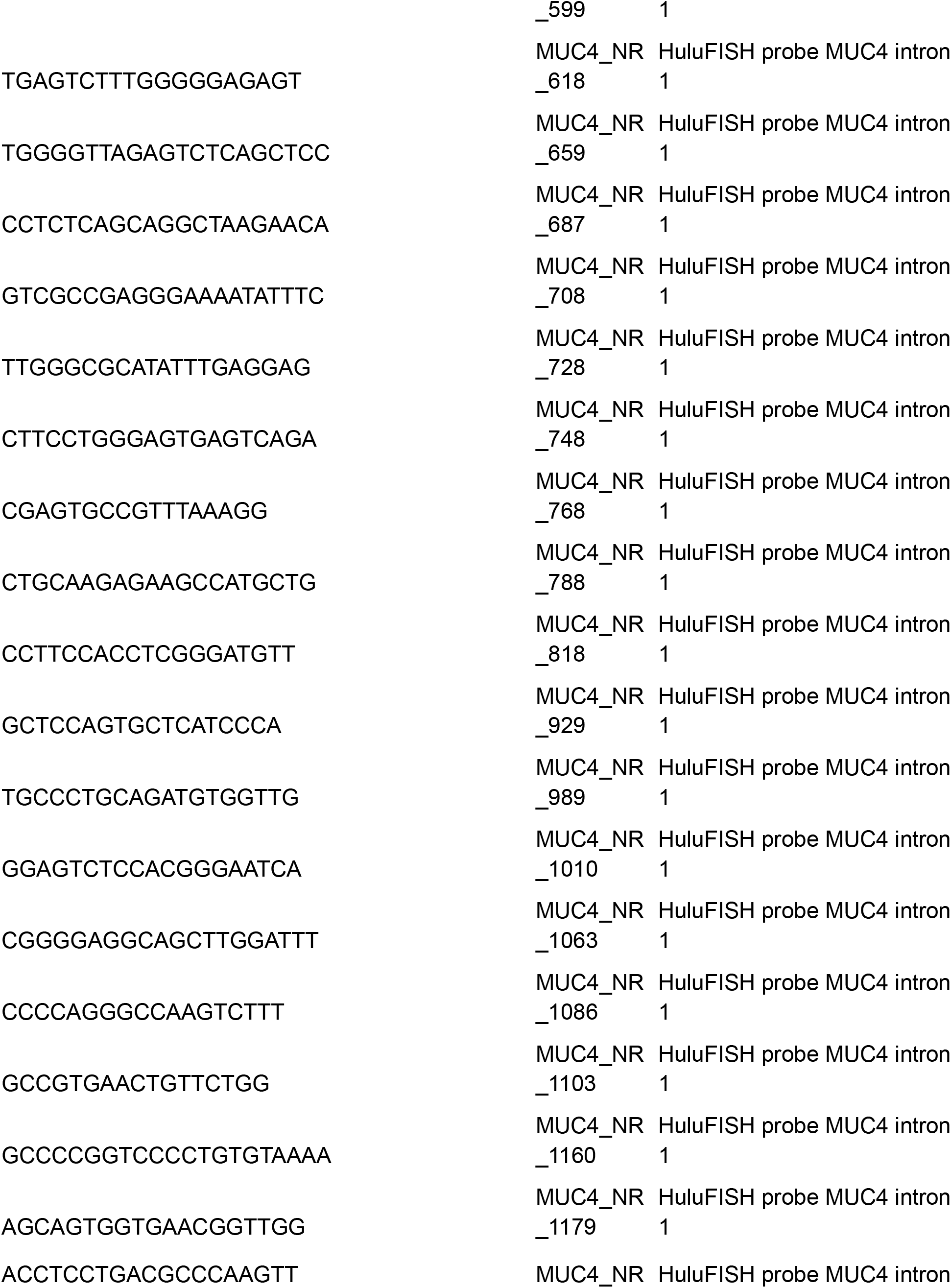
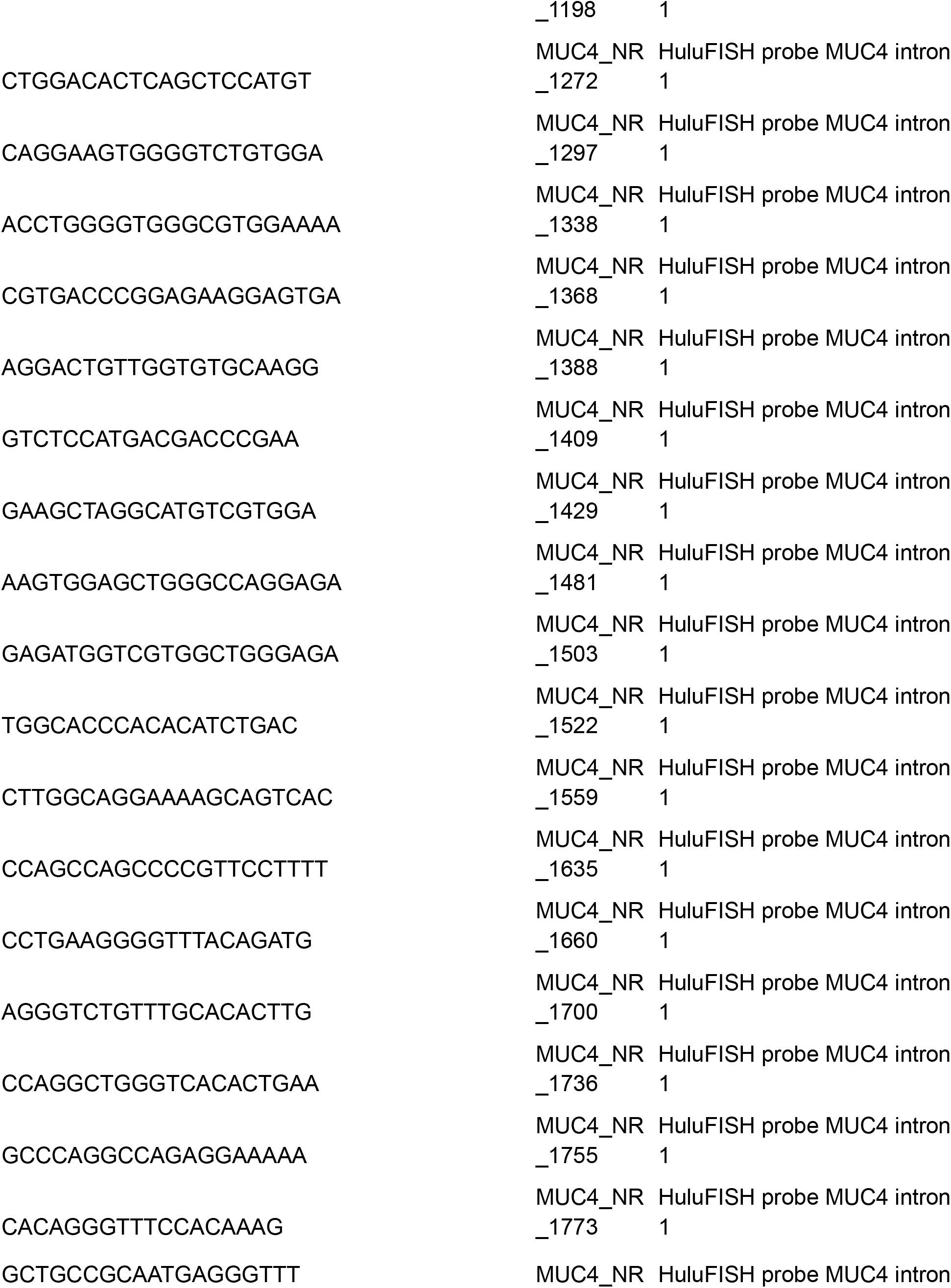
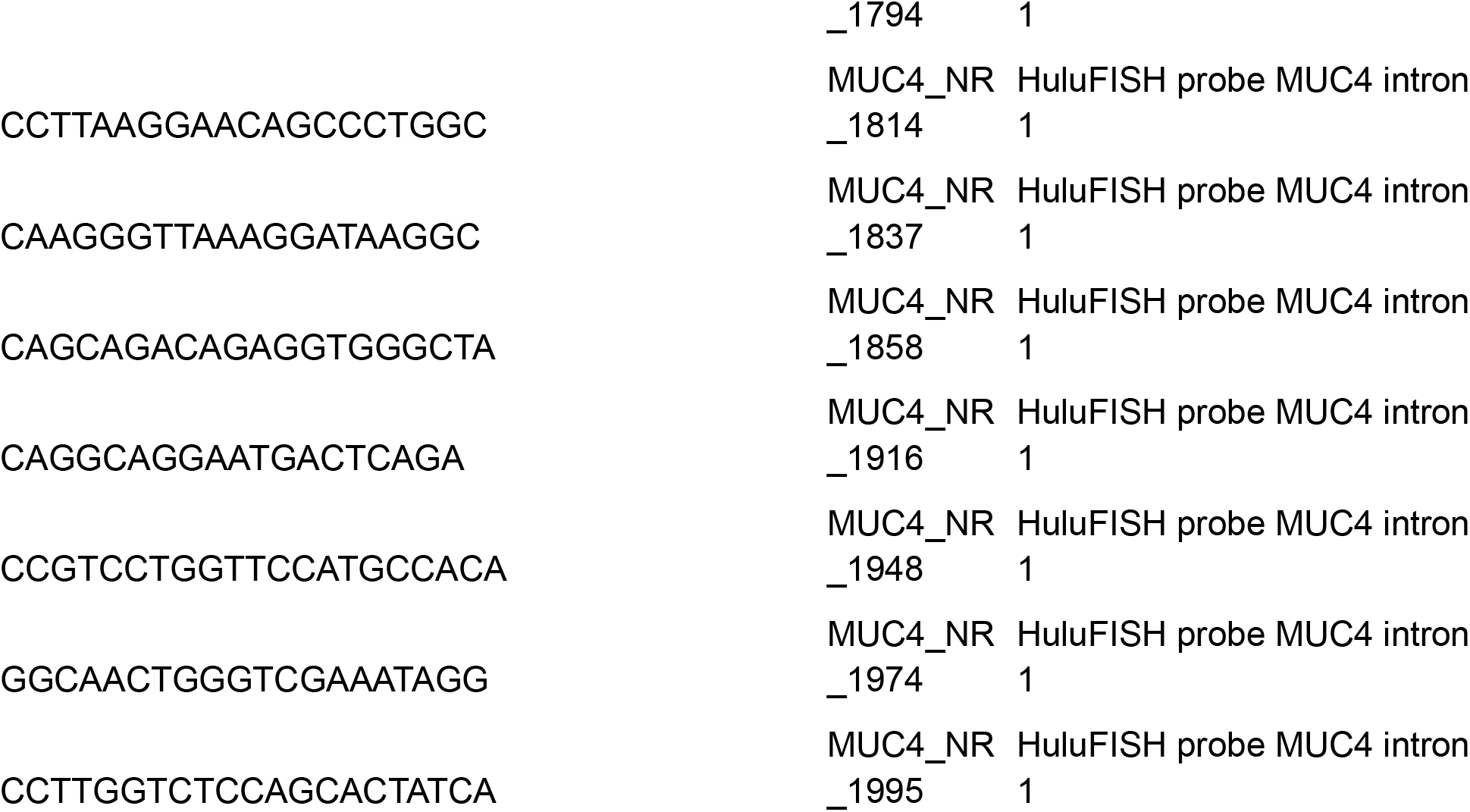
Sequence of oligos used in this study.

